# Site-specific K63 ubiquitinomics reveals post-initiation regulation of ribosomes under oxidative stress

**DOI:** 10.1101/392076

**Authors:** Songhee Back, Christine Vogel, Gustavo M. Silva

## Abstract

During oxidative stress, K63-linked polyubiquitin chains accumulate in the cell and modify a variety of proteins including ribosomes. Knowledge of the precise sites of K63 ubiquitin is key to understanding its function during the response to stress. To identify the sites of K63 ubiquitin, we developed a new mass-spectrometry based method that quantified >1,100 K63 ubiquitination sites in yeast responding to oxidative stress induced by H_2_O_2_. We determined that under stress, K63 ubiquitin modified proteins involved in several cellular functions including ion transport, protein trafficking, and translation. The most abundant ubiquitination sites localized to the head of the 40S subunit of the ribosome, modified assembled polysomes, and affected the binding of translation factors. The results suggested a new pathway of post-initiation control of translation during oxidative stress and illustrated the importance of high-resolution mapping of noncanonical ubiquitination events.

## Introduction

Ubiquitination is one of the most prevalent post-translational modifications in eukaryotes, modifying thousands of proteins and involving hundreds of enzymes.^1–3^ It is primarily known as the signal for protein degradation in the nucleus and the cytosol.^3^ However, a variety of ubiquitin chain types can be conjugated to a given substrate, and their functions expand to protein trafficking, protein-protein interaction, and other signaling processes.^4,5^ This diversity in ubiquitin chain types is determined by the internal lysine residues linking the ubiquitin molecules. K48 and K11 ubiquitin chains are the main signal for protein degradation while K63 ubiquitin is the only linkage not related to protein degradation by the proteasome.^4,6–8^

The abundance and the dynamics of the different chains and signals can vary according to the cellular environment. In response to oxidative stress, multiple ubiquitin chain types rapidly and strongly accumulate in cells including those with K63 and K48 linkages.^9^ We have recently shown that K63 polyubiquitin chains modified ribosomes, retained polysomes, and supported cellular survival during oxidative stress.^9^ However, the exact sites of K63 ubiquitin and its role in ribosome function remained unknown. However, knowing the modification sites is a crucial next step to understand how ubiquitin regulates ribosome activity and protein expression.

Given the multitude of different ubiquitination types that affect the ribosome and other proteins,^10–12^ we need tools to identify, map, and disentangle the distinct ubiquitin signals to understand their physiological role *in vivo.* Mass spectrometry is the current tool of choice to investigate protein ubiquitination and its physiological roles.^13^ Mass spectrometry analysis typically identifies ubiquitinated lysines via the di-glycyl remnant (GG) left by tryptic digest of the target protein^,14,17-23^ Importantly though, this tryptic digest also processes the ubiquitin chain, removing information of the linkage type that modified the protein.

Therefore, we developed a new method, called Ub-DiGGer, which comprises a sequential enrichment strategy that preserves information on the linkage type, in this case K63, but also extracts modification sites with high specificity and proteome coverage. Using Ub-DiGGer, we identified ~1,100 K63 ubiquitination sites in the yeast proteome responding to stress. The modification affected several pathways, such as translation, amino acid transport, and the Golgi apparatus. We investigated the modification of the ribosome in more detail and present results that suggest a role of K63 ubiquitin in translation shutdown upon oxidative stress in a mechanism independent of the canonical inhibition of initiation via reduced availability of the ternary complex.

## Materials and Methods

### Yeast strains and growth

All yeast *Saccharomyces cerevisiae* strains used in this study are described in Supplementary Table S3. Standard recombination methods were used to delete and tag genes, which were confirmed by PCR and western blot, respectively.

### Sensitivity assay

Cells were grown to logarithmic phase in minimal medium containing 0.67% yeast nitrogen base, 2% dextrose, and required amino acids at 30°C. Cellular density was normalized to OD_600_ of 0.2. Cells were then sequentially diluted at 1:5 ratio and plated in YPD rich medium plates containing: 50 ng/ml rapamycin, 20 ug/ul anisomycin, 0.6 or 1.2 mM H_2_O_2_.

### Isolation of K63 ubiquitinated peptides

The sequential ubiquitin enrichment was applied to identify and quantify the lysine residues modified by K63 ubiquitin in response to oxidative stress. For each biological replicate, 2.4 liters of each strain (WT and K63R) were cultured to mid-log phase into synthetic SILAC media supplemented with light and heavy isotopes of lysine and arginine, respectively (L-Arg6 ^13^C; L-Lys8 ^13^C, ^15^N; Cambridge Isotopes). Cells were grown for at least 10 generations in the SILAC medium before being treated with 0.6 mM H_2_O_2_ for 30 min at 30 °C. Equal amounts of WT and K63R cells were collected by centrifugation, mixed, and disrupted by glass-bead agitation at 4 °C in lysis buffer: 50 mM Tris-HCl pH 7.5, 100 mM NaCl, 5 mM EDTA, 20 mM chloroacetamide, 50 nM FLAG K63-TUBE peptide (LifeSensors), 1X EMD Millipore protease inhibitor cocktail set I. Approximately 350 mg of protein were loaded onto a column containing M2 anti-FLAG resin (Sigma). Sample was eluted with 0.1 M glycine pH 2.5, neutralized with NaOH and buffered with 100 mM Tris-HCl pH 8.0, 1 mM CaCl_2_ prior to overnight digestion with Trypsin/Lys-C (Promega). Samples were desalted using C18 Hypersep Spin Column (Thermo), dried completely and resuspended in IAP buffer from PTMScan Ubiquitin K-ε-GG kit (Cell Signaling). For the enrichment of diglycyl-lysine peptides (GG), tryptic peptides were incubated for 4h at 4 °C with anti-GG beads and eluted with 0.15% formic acid according to the manufacturer’s protocol. Samples were desalted once again with Hypersep tips and resuspended in 0.1 % FA, prior to LC-MS/MS analysis.

### Polysome proteomics

For the quantitative analysis of the composition of polysomes and monosomes, two biological replicates were prepared from cells with and without 30 min of 0.6 mM H_2_O_2_ treatment. Cell disruption was performed as described below and approximately 400 μg of RNA from each yeast strain wild-type and K63R grown in heavy and light SILAC medium, respectively, were sedimented in a sucrose gradient. The monosome fraction (40S, 60S and 80S) was separated from the polysome fractions, proteins were precipitated with 10% trichloroacetic acid (TCA), protein pellet was washed twice with ice-cold acetone, dried out, and resuspended in 8M urea buffered in 50 mM Tris-HCl pH 8.0, 1 mM CaCl_2_. Samples were digested in a two-step fashion by incubating with Trypsin/Lys-C mix (Promega) for 3h followed by dilution of urea to 1M with Tris-buffer pH 8.0 for overnight tryptic digestion at 37 °C. Samples were desalted using Hypersep spin tips (Thermo) prior to LC-MS/MS analysis.

### Mass spectrometry analysis

Protein preparation for proteomics analysis was performed as previously described^9^. Briefly, tryptically digested proteins isolated by polysome profiling were separated on a 25-cm Thermo Acclaim PepMap C18 column (75 μm ID, 2 μm particle size, 100 Å pore size, cat # 164734) by reverse-phase chromatography using a gradient of 5–60% acetonitrile over 2.5 h, performed on an Eksigent NanoLC 2DPlus liquid chromatography system. The eluted peptides were injected in-line into an LTQ Orbitrap Elite mass spectrometer (Thermo Scientific). Data-dependent analysis was performed at a resolution of 120,000 at the MS level, AGC at 1e6, maximum injection time of 100 ms, with the top 20 most intense ions selected from each MS full scan, and dynamic exclusion set to 30 seconds if m/z acquisition was repeated within a 30-second interval. MS2 was acquired at a resolution of 15,000, AGC at 5e4 and maximum injection time of 100ms. The K63 ubiquitinomics samples were loaded onto a 50 cm Easy Spray PepMap C18 column (75 μm ID, 2 μm particle, 100 Å pore size, cat. No ES803) in-line with a Q-Exactive (Thermo Scientific) mass spectrometer using a 75-min gradient (0-40% ACN). Data dependent analysis was performed at resolution of 70,000 at the MS level, AGC at 1e6 and maximum injection time of 120 ms. Top 20 most intense ions selected from each MS full scan, with dynamic exclusion set to 30-second if m/z acquisition was repeated within a 30-second interval. MS2 was acquired at a resolution of 17,500, AGC at 5e4 and maximum injection time of 120ms. The RAW data files were processed using MaxQuant to identify and quantify protein and peptide abundances. The spectra were matched against the yeast *Saccharomyces cerevisiae* Uniprot database. Protein identification was performed using 10 ppm tolerance with a posterior global FDR of 1% based on the reverse sequence of the yeast FASTA file. Up to two missed trypsin cleavages were allowed, and oxidation of methionine and N-terminal acetylation were searched as variable post-translational modification and cysteine carbamidomethylation as fixed. For K63 ubiquitinated peptides, diglycyl-lysine (GlyGly(K)) was also searched as variable modification.

### Polysome analysis

Polysome profiling was performed as described previously^9^. Briefly, a total of 400 μg of RNA was sedimented by ultracentrifugation for 150 min at 38,000 rpm (Beckman SW41Ti rotor) at 4 °C. MgCl_2_ concentration varied from 1 to 3 mM in the lysis buffer and from 1 to 30 mM in the sucrose buffer. Polysome traces were collected with PeakChart software coupled to a Density Gradient Fractionation System (Brandel).

### Western blot

Proteins were separated by standard 10% SDS-PAGE, with samples loaded in Laemmli buffer, transferred for 2h to a PVDF membrane, and immunoblotted using the following antibodies: anti-K63 ubiquitin (1:4,000; EMD Millipore, cat. No. 05-1308, clone apu3); anti-GAPDH (1:4,000; Abcam, cat. No. ab9485); anti-Rps10 (1:6,000, Sigma, cat no. WH0004736M1); anti-Rps6 (1:4,000; Abcam cat no. ab40820), anti-puromycin (1,2:500, EMD Millipore MABE343, clone 12D10), anti HA (1:2,500; Invitrogen, cat no. 71-5500), anti-K48 ubiquitin (1:8,000; Cell Signaling, cat. No 8081).

### Translation activity

Puromycin incorporation into newly synthesized proteins was performed to measure translation activity. Cells were treated in the presence or absence of 0.6 mM H_2_O_2_ for 15 min prior to 30 min incubation with 0.9 mM puromycin. Cells were collected by centrifugation and lysate was prepared as described above. Twenty micrograms of protein were loaded onto a 10% SDS-PAGE gel for immunoblotting.

## Results

### In-depth characterization of the K63 ubiquitin-modified yeast proteome

We developed a mass spectrometry-based method that maps both, the sites of ubiquitin modification for a particular linkage type (Figure 1A). The method, called Ub-DiGGer, first isolates the targets of a selective linkage with a highly specific enrichment tool. The isolated proteins are then digested with trypsin and subjected to a second enrichment step that isolates the di-glycyl (GG) remnant on the modified lysine. Finally, the isolated peptides are identified by mass spectrometry (Figure S1A). To quantify K63 ubiquitin changes in response to oxidative stress and to monitor the specificity of the method, we combined Ub-DiGGer with Stable Isotopic Labeling in Cell culture (SILAC)^14^ of wild-type cells compared to the K63R ubiquitin mutant, which is incapable of assembling K63 ubiquitin chains (Figure S1B). Using the K63R ubiquitin mutant strain as a control to account for false positive identifications, we mapped 1,113 K63 ubiquitin sites at 1% peptide based false discovery rate.

**Figure 1.**
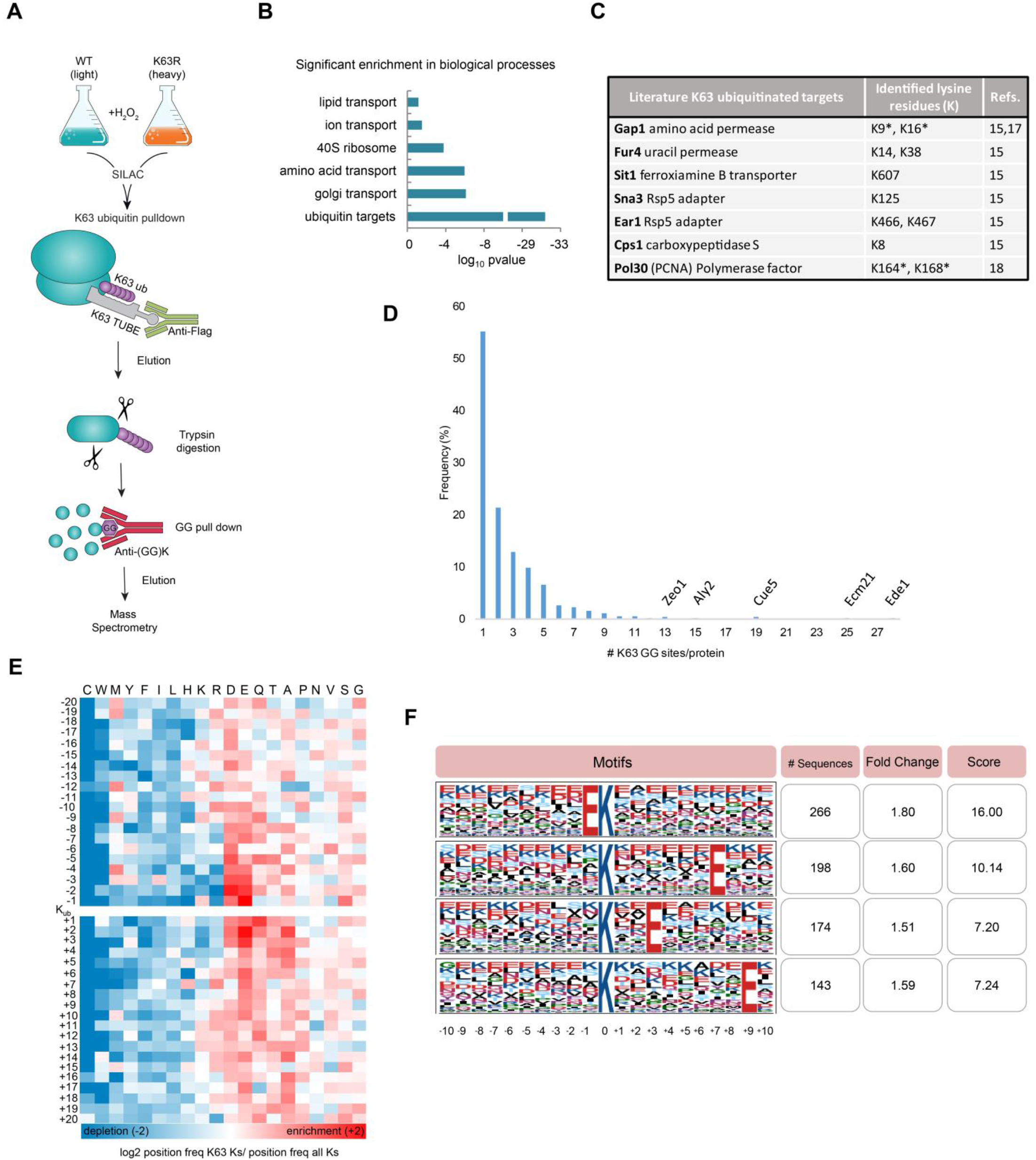
K63 ubiquitin-modified proteome in response to oxidative stress. **(a)** Scheme describing the Ub-DiGGer method to identify and quantify K63 ubiquitinated lysine residues. **(b)** Histogram of Gene Ontology function annotations for K63 target proteins using DAVID functional annotation tool. *p-values* are Bonferroni-corrected for multiple hypotheses testing. **(c)** Sites identified for known K63 ubiquitinated proteins. *K63 ubiquitin sites previously identified and confirmed by this study. **(d)** Frequency distribution of number of K63 GG sites per protein. **(e)** Heatmap shows log base 2 ratio of the frequency of the amino acid position relative to the K63 ubiquitin modified lysine (K_ub_) over the frequency of the amino acids at the respective position on all lysine residues in the yeast proteome. The order of the amino acids (columns) was based on hierarchical clustering of their ratio profiles. Fold change enrichment is shown in red and depletion is shown in blue. **(f)** K63 ubiquitin motif analysis determined by Motif-X^40^. Top four score motifs are displayed centered on the ubiquitinated lysine and a window of ± 10 residues.

We selected K63 ubiquitinated peptides whose SILAC ratios changed at least 1.6-fold in response to stress, consistently across both replicates illustrating the high quality and reproducibility of the data (Table S1, Figure S1A). The peptides mapped to 490 distinct proteins (Table S1).

The identified peptides showed a significant enrichment for ubiquitinated proteins (Figure 1B, p-value <e-30) further validating our approach. In addition, the peptides arose from proteins from the Golgi apparatus and amino acid transport (p-value < e-6, Figure 1B). Other targets functioned in autophagy, ion transport, carbohydrate metabolism, endocytosis, and nutritional response (Figure S1C). These results were in agreement with reports that many proteins involved in trafficking, metabolism, and translation are regulated by K63 ubiquitin^15, 16^.

To-date, a handful of K63 targets and sites have been confirmed for yeast, which we used for further validation of our dataset. Indeed, our dataset reported K63 ubiquitin sites for the permeases Fur4 (K14, K38) and Gap1 (K9, K16), the transporter Sit1 (K607), the Rsp5 adapters Sna3 (K125) and Ear1 (K466, K467) and for the carboxypeptidase Cps1 (K8). All these proteins have been previously characterized as targets of K63 ubiquitin^15^. Moreover, our data retrieved the known K63 modified lysine residues in Gap1, i.e. K9 and K16^17^. We also identified K164 in PCNA, a well-characterized site of K63 ubiquitin important for DNA repair signaling^18^. Moreover, we found that most targets contained three or fewer K63 ubiquitin sites (Figure 1D), but some proteins involved in endocytosis or cell wall organization were highly decorated with K63 ubiquitin chains, having more than ten GG sites (*e,g*, Ede1, Ecm21, and Zeo1, Figure 1D).

Our results also corroborated the hypothesis of the absence of a sequence motif signaling for ubiquitination^19^. Importantly, our data arose from a very specific subset of ubiquitination events: K63 ubiquitin changes in response to oxidative stress. Despite this highly focused dataset, we did not identify a sequence motif surrounding the K63 ubiquitin sites (Figure 1E and 1F). However, we identified physicochemical properties surrounding the modified sites: i) an enrichment of acidic amino acids (glutamate and aspartate); ii) an enrichment in glutamine and alanine in the positions immediately following the modified lysine; and iii) a major depletion of cysteine and tryptophan surrounding the modified lysine (Figure 1E). Therefore, our results support the school of thought that ubiquitination does not occur exclusively via a recognition motif^19^. Instead, structural components associated with amino acid sequence might be recognized by ubiquitination enzymes.

Many of the K63 ubiquitination events in response to stress mapped to the ribosome, and we investigated these events in more detail. These sites were significantly enriched in components of the 40S ribosome (Figure 1B, p-value < e-3.5). In total, 78 K63 ubiquitin sites mapped to 37 ribosomal proteins, and amongst them, 45 sites mapped to 40S proteins and three additional sites mapped to Asc1 (RACK1), a 40S-associated protein (Table S1). This preference is substantial given that the 40S subunit is much smaller and has fewer lysine residues than the 60S (539 and 834, respectively). Further, the SILAC ratios for the 40S subunit sites were significantly higher than those for the 60S (Figure S1D, t-test, p < e-6).

When mapping K63 ubiquitin sites to a publicly available X-ray structure of the yeast 80S ribosome^20^, we found that most sites were located at the solvent exposed surface of the ribosomes (Figure 2A). Moreover, the majority of sites, including those that changed most drastically under stress, clustered at the head of the 40S subunit (Figure 2 and Figure S1E).

**Figure 2.**
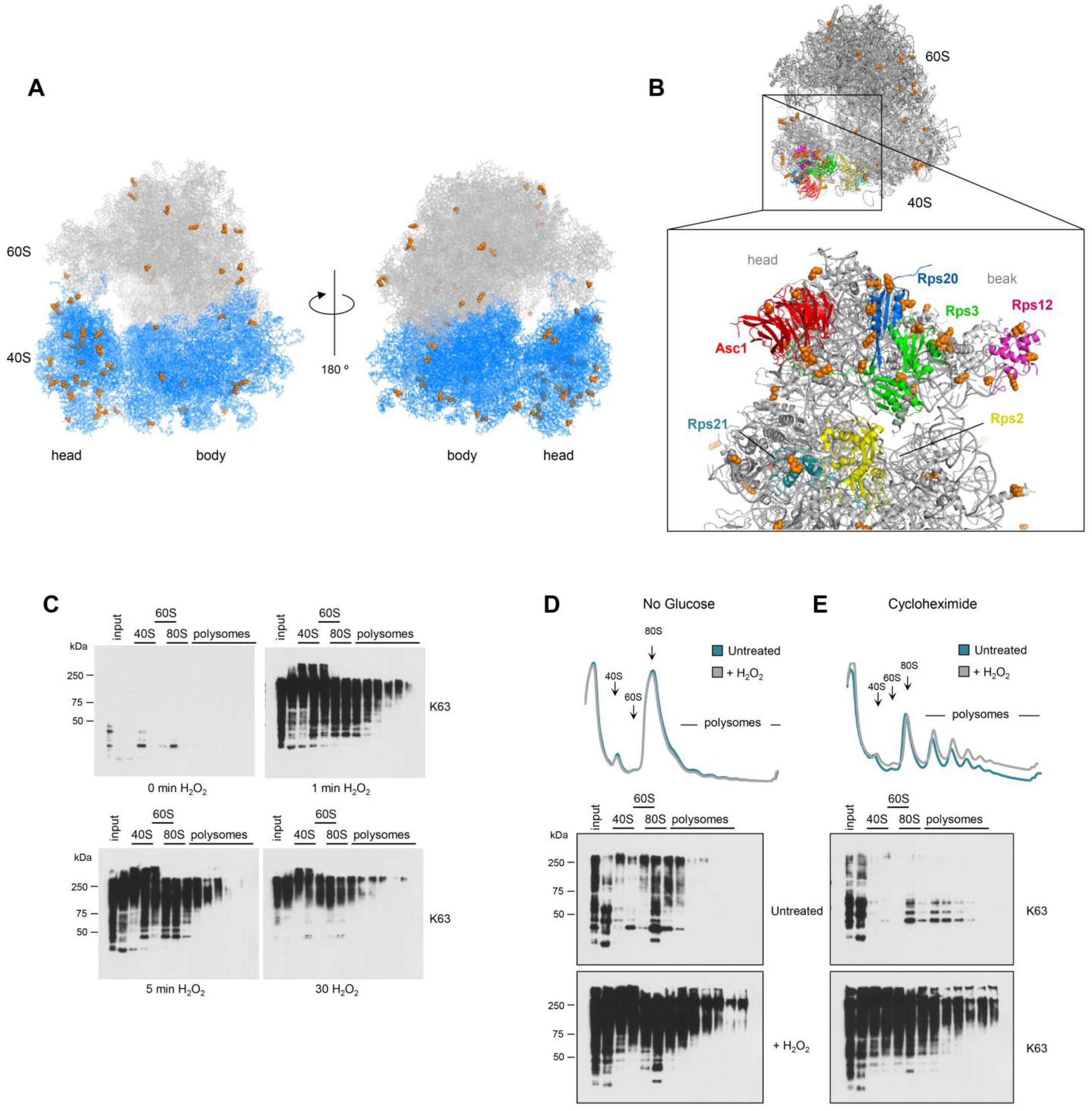
K63 ubiquitin modifies the head of the 40S ribosome. Mapping of the K63 ubiquitinated sites (orange) to the 3D structure of the yeast ribosome (PDB id:4v88)^20^. Figures highlight K63 modification at **(a)** the 80S ribosome and **(b)** at the head of the ribosome showing the most abundantly K63 ubiquitinated subunits colored in yellow (Rps2/uS5), green (Rps3/uS3), magenta (Rps12/eS12), blue (Rps20/uS10), and teal (Rps21/eS21). Ribosome associated protein Asc1 (RACK1) is shown in red. **(c)** Anti-K63 ubiquitin western blot of ribosome fractions from cells treated with H_2_O_2_ for the designated times. **(d,e)** Sucrose sedimentation profiles of polysome from wild-type cells in which translation was inhibited **(d)** at initiation after cells were transferred to glucose depleted medium or **(e)** at elongation after treatment with 150 μg/ml cycloheximide. Anti-K63 ubiquitin blot was performed for cells cultivated in the presence or absence of H_2_O_2_. WT, wild-type SUB280 yeast strain. K63R, ubiquitin K63R mutant SUB413 yeast strain.

### K63 ubiquitin regulates translation post initiation

First, we showed that K63 ubiquitin rapidly affected free subunits (40S and 60S), monosomes (80S), and polysomes after stress induction by H_2_O_2_ (Figure 2C). We tested if modification of assembled 80S ribosomes arose from the unassembled subunits being modified prior to their assembly or if K63 ubiquitin affected mono- and polysomes directly. We assayed for this question measuring K63 ubiquitination of polysomes in which translation was inhibited and therefore no assembly or progression of monosomes occurred. Using glucose deprivation or cycloheximide to inhibit initiation and elongation, respectively, K63 ubiquitin modified both free subunits and mono- and polysomes (Figure 2D and S2E), supporting our hypothesis that K63 ubiquitin directly targets assembled ribosomes and polysomes.

The entire head of the 40S ribosome is a flexible and dynamic structure that harbors the decoding center, surrounds the mRNA entry and exit site, engages tRNAs, and is heavily regulated during the different stages of translation^21, 22^. Therefore, we conducted additional experiments to test for the impact of the extensive K63 ubiquitination of the 40S subunit on ribosome activity during oxidative stress.

First, we tested for the specific stage of translation that was affected by K63 ubiquitination. During oxidative stress, translation initiation is rapidly inhibited by the Gcn2 kinase that phosphorylates the initiation factor eIF2*α*, thereby reducing the levels of translation-competent ribosomes^23, 24^. To test whether K63 ubiquitin interfered with inhibition of translation initiation by Gcn2, we analyzed polysome levels by sucrose sedimentation profiling. The K63R ubiquitin mutant strain displayed lower levels of polysomes when compared to the wild-type under stress (Figure 3A and 3B). However, both strains maintained the polysome levels when *GCN2* was deleted and initiation was fully operational under stress (Figure 3C and 3D). This finding suggested that K63 ubiquitin’s function was independent of the inhibition of initiation. Supporting this conclusion, cells deleted for *GCN2* in the K63R background *(K63R-gcn2Δ)* did not rescue the K63R cell’s sensitivity to translation inhibitors (Figure S2A). Second, we showed that ribosomes for the K63R strain are less stable than the wild-type. At low MgCl_2_ concentration, which destabilizes 80S particles, samples from K63R cells under stress were devoid of both mono- and polysomes and consisted mostly of free 40S and 60S subunits. In contrast, 80S, disomes, and trisomes were still observed in the wild-type strain under stress (Figure 3E).

**Figure 3.**
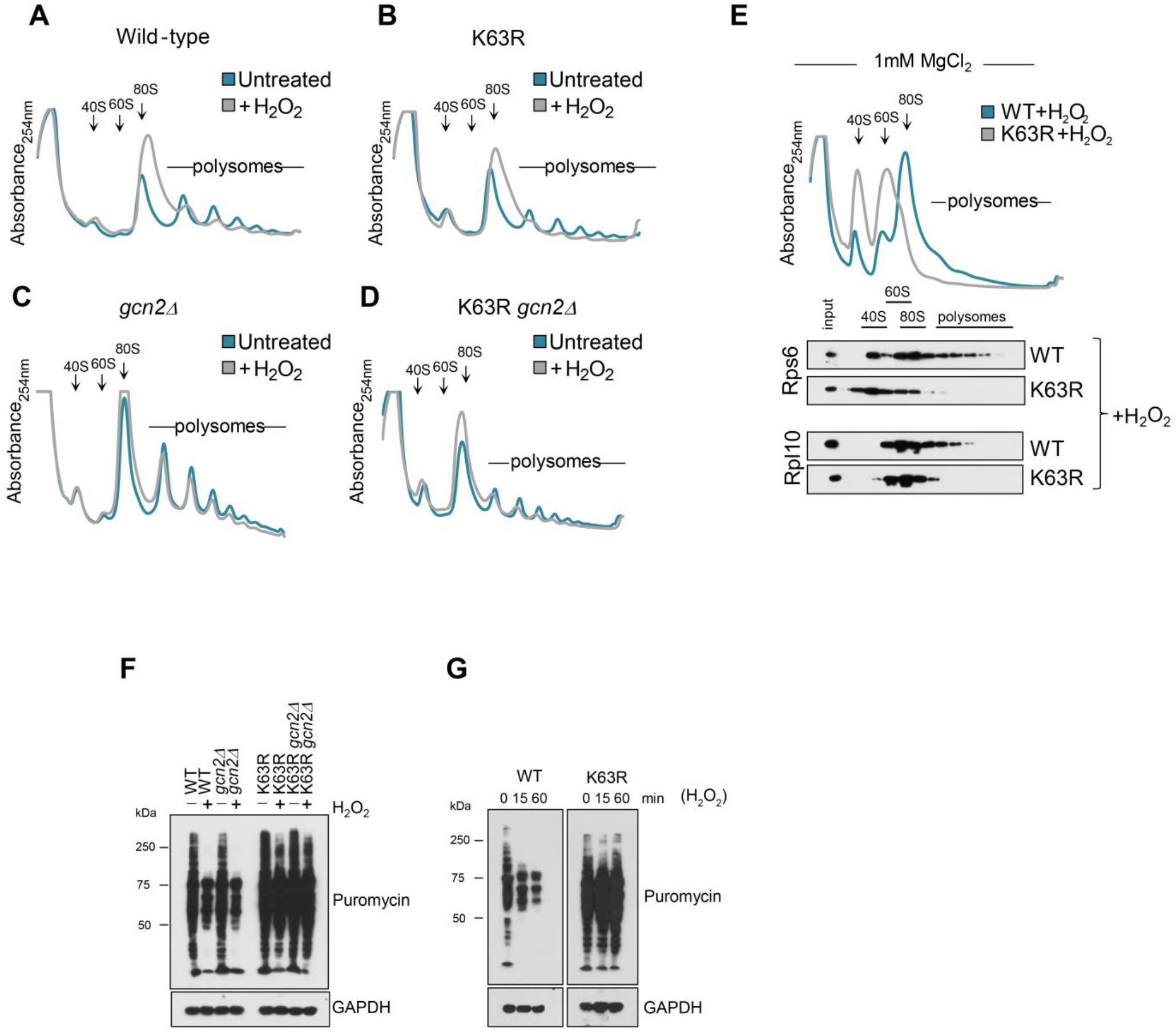
K63R ribosomes are more sensitive to stress than wild-type, independently of *GCN2.* Sucrose sedimentation profiles of polysome from **(a)** wild-type, **(b)** K63R, **(c)** *gcn2Δ,* and **(d)** K63R-gcn2Δ Cells were incubated in the presence or the absence of 0.6 mM H_2_O_2_ for 30 min. Anti-K63 ubiquitin blots of the same experiment are present in Figure S2B. **(e)** Polysomes from wild-type and K63R under H_2_O_2_ were extracted at low magnesium chloride concentration (1 mM). Western blot of sedimentation fractions was performed for 60S large and 40S small subunit proteins Rpl10 and Rps6, respectively. WT, wild-type SUB280 yeast strain. K63R, ubiquitin K63R mutant SUB413 yeast strain. **(f,g)** Western blot against puromycin was used to evaluate *de novo* protein synthesis. Cells were treated with H_2_O_2_ for 15 or 60 min prior to treatment with 0.9 mM puromycin for additional 30 min. Anti-GAPDH blot was used as loading control.

Considering the 40S location of the K63 ubiquitin sites, the low stability of K63R ribosomes, and the independence from translation initiation, our data suggested that K63 ubiquitin might regulate ribosome activity *post* initiation. Inhibition of translation at elongation had been proposed during oxidative stress but without a known mechanism^23–26^. To test for post-initiation regulation mediated by K63 ubiquitin, we measured translation output by monitoring the incorporation of puromycin into newly synthesized proteins. The wild-type cells showed low puromycin incorporation in response to H_2_O_2_, confirming strong translation repression (Figure 3F). In agreement with earlier findings^23, 26^, deletion of the *GCN2* was insufficient to restore translation output under stress, suggesting additional inhibition of translation beyond initiation (Figure 3F). In the absence of stress, K63R mutant cells had similar puromycin incorporation as the wild-type indicating normal translation levels. However, translation output remained unaltered in the K63R mutant in response to H_2_O_2_ suggesting that K63 ubiquitin was involved in translation halt, independently of initiation (Figure 3F).

In the K63R mutant, normal translation output could be achieved either by constant production of protein that is unaffected by the stress or by elongating ribosomes that remain on the mRNAs and still translate during the early phases of the stress response. In the latter scenario, we would expect puromycin incorporation to decrease over time, as there would be little to no *de novo* translation initiation due to Gcn2 activity during stress. To test this hypothesis, we used short (15min) and long (1h) H_2_O_2_ incubation and monitored protein synthesis. We found that incorporation of puromycin occurred at normal levels in the K63R mutant compared to the repressed wild-type regardless of the duration of the stress (Figure 3G). These findings provided another line of evidence for K63 ubiquitin acting via a mechanism independent of the inhibition of translation initiation.

### K63 ubiquitin alters the ribosome’s interaction with translation factors

K63 ubiquitin can act as a scaffold, recruit ubiquitin-binding proteins, or prevent binding of additional proteins, thus affecting a variety of signaling pathways independently of proteasomal degradation^6^. Since our results indicated that K63 ubiquitin modifies assembled monosomes and polysomes (Figure 2C-2E), and affects their stability and activity (Figure 3), we tested if these roles arose from K63 ubiquitin impacting the binding of regulatory factors to the ribosome.

We examined two candidate factors which are known to regulate ribosome function: the ribosome associated factor Asc1 (RACK1)^11^ and the E3 ubiquitin enzyme Hel2^12^. Asc1 (RACK1) was modified by K63 ubiquitin in our experimental system (Supplementary Table S1), and localized next to K63 ubiquitin sites on the 40S ribosome (Figure 2B). We therefore tested the effect of *ASC1* deletion on ribosome ubiquitination, stability, and stress response. Although deletion of *ASC1* rendered yeast cells more sensitive to oxidative stress and translation inhibitors (Figure S3A), *ASC1* deletion did not result in active translation (Figure S3B), reduced abundance of polysomes (Figure S3C), or diminished K63 ubiquitin levels (Figure S3C) – the molecular phenotypes observed in the K63R mutant strain. ASC1 therefore appeared to work independently of K63 ubiquitination. Similarly, we showed that the ubiquitin ligase Hel2 was not responsible for the accumulation of K63 ubiquitin in response to oxidative stress (Figure S3D).

To systematically screen for other factors whose binding to the ribosome could be affected by K63 ubiquitin during oxidative stress, we used mass spectrometry to analyze samples from the sucrose sedimentation profile (Figure 4A, Figure S4A, Supplementary Table S2). While the abundance of ribosomal proteins was mostly unchanged, many translation factors that were more abundant in the wild-type compared to the K63R mutant. This effect was observed in the monosomal (Figure 4B) and also in the polysomal fraction (Figure S4B). Amongst these factors, we identified several components of the eukaryotic initiation factor eIF3 complex, e.g. Nip1 (eIF3a) and Tif35 (eIF3g)(Figure 4B, Figure S4B). eIF3 is a key translation factor which binds to the 40S subunit, stabilizes the ternary complex, recruits capped mRNA, and assists translation initiation^27, 28^.

**Figure 4.**
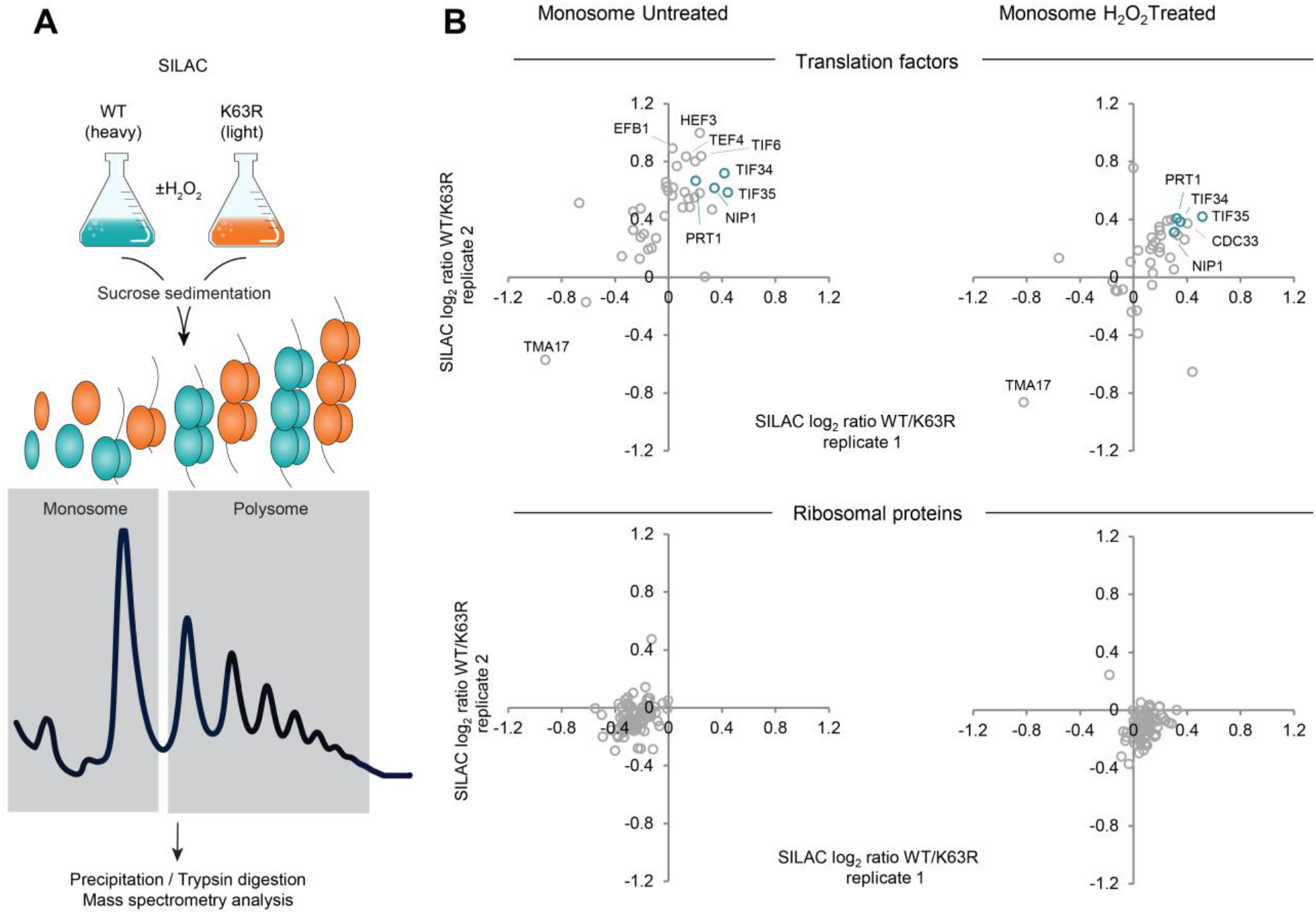
Ribosome proteomics reveals K63 ubiquitin-dependent binding of translation factors. **(a)** Scheme of polysome proteomics method following sucrose sedimentation profiling. **(b)** Scatter plot of log_2_ SILAC WT/K63R ratios of protein intensity from two independent replicates in the monosome fraction. *(top)* SILAC ratio of translation factors, *(bottom)* SILAC ratio of ribosomal proteins. Subunits of the eIF3 translation initiation factor are labeled in teal. WT, wild-type GMS280 yeast strain. K63R, ubiquitin K63R mutant GMS413 yeast strain

We tested for both Nip1 and Tif35 if their binding to the ribosome depended on K63 ubiquitin. While the abundance of Nip1 bound to the ribosome remained similar in both the wild-type and the K63R strain (Figure S5A), Tif35’s abundance decreased in the K63R ubiquitin mutant strain compared to wild-type (Figure 5A and 5B). These results were robust to changes in blotting exposure and sample loading (Figure S5B and S5C). Consistently, reduced expression of *TIF35* in the DAmP strain^29^ sensitized cells to oxidative stress and to translation inhibitors (Figure S6A), and slightly increased translation as measured by puromycin incorporation (Figure S5A). Tif35 association with the ribosome might therefore be affected by K63 ubiquitin.

**Figure 5.**
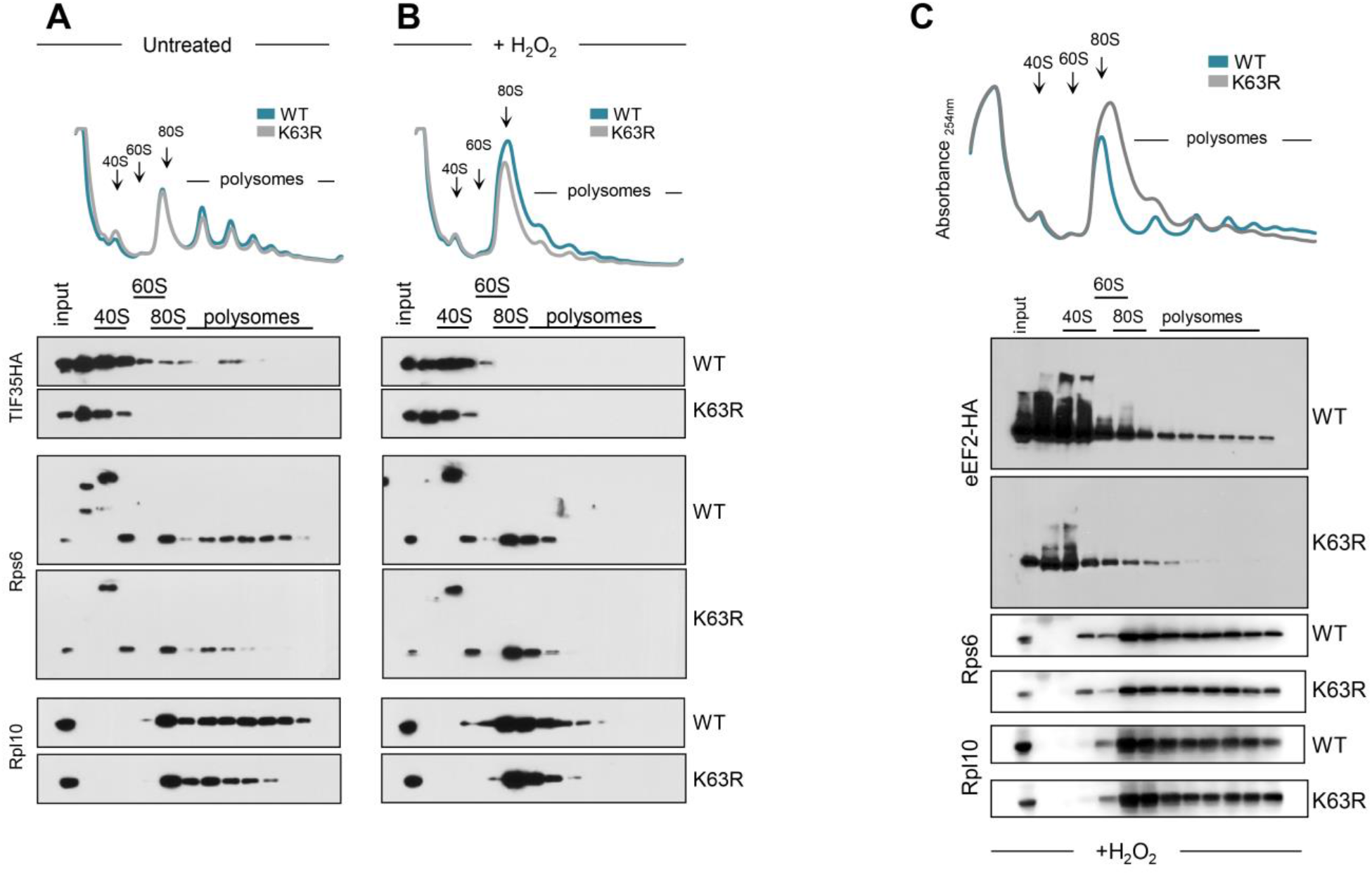
Tif35 and eEF2 are depleted in K63R ubiquitin polysomes. Sucrose sedimentation profiles of polysomes from **(a)** wild-type and **(b)** K63R mutant strain, expressing HA-tagged Tif35. (c) Sucrose sedimentation profiles of polysomes from wild-type and K63R mutant strain, expressing HA-tagged eEF2. Cells were incubated in the presence or absence of 0.6 mM H_2_O_2_ for 30 min. Western blot of sedimentation fractions were performed for anti-HA, 60S large protein Rpl10, and 40S small protein Rps6.

Finally, we demonstrated that the K63R ubiquitin mutation also affected the binding of the elongation factor eEF2, but not eEF3. eEF2 is the elongation factor that binds to the ribosome during the translocation stage of elongation^30^, while eEF3 is a yeast-specific elongation factor involved in the post-translocation step of translation, aiding the release of tRNAs from the E-site^31^. We found that in wild-type cells, a larger amount of eEF2 was present and proportionally bound to the ribosome compared to the K63R mutant (Figure 6D). In comparison, eEF3 binding did not change (Figure S6C). Our results suggested that the accumulation of K63 ubiquitin on the 40S ribosome affected the interaction of the particle with translation factors, ultimately hindering the progression of the elongation cycle in response to oxidative stress.

## Conclusions

We developed a new mass spectrometry-based method, Ub-DiGGer, to monitor the K63 ubiquitin-modified proteome in response to oxidative stress. The novelty of the method lies in its ability to identify the targets of a specific ubiquitin linkage (K63) and the modified lysine residues simultaneously. Previous methods successfully characterized either the ubiquitin modified proteome^19, 32^ or quantified the cellular abundances of different ubiquitin linkages^7, 33^.

Our method selectively isolated the K63 ubiquitinated proteins prior to tryptic digest and hence enabled large-scale and quantitative site analysis (Figure 1A, Table S1). The data provided in this study is unbiased, large scale, quantitative, linkage-specific, and was acquired in response to defined conditions – all aspects that render the information highly useful for biological interpretation. Given the emergence of new enrichment methods of sufficient sensitivity and specificity, Ub-DiGGer can be expanded to other ubiquitin linkage types.

Our dataset of >1,100 sites provided the largest resource on K63 ubiquitination to date and will foster our understanding of K63 ubiquitin targets and functions in the cell. K63 ubiquitin targeted proteins from Golgi-mediated, ion, and amino acid transport, endocytosis, and carbohydrate and lipid metabolism (Table S1). We illustrated the wealth in information gained from our data at the example of translation control. Finally we showed that our method was able to retrieve proteins and ubiquitin sites that were previously confirmed as K63 ubiquitin substrates^15, 17^ supporting the importance and validity of the approach.

We further illustrated the significance of this dataset by investigating the functional role of K63 ubiquitin in translation regulation. We first showed that a large cluster of K63 modified sites are located at the head of the 40S subunit (Figure 2), which is ribosome’s regulatory center^21, 22, 28^. K63 ubiquitin acts independently of *GCN2* and therefore, independently of initiation control (Figure 3C, 3D and 3F). Although translation initiation is largely inhibited under oxidative stress, it has been shown that elongation transit time and ribosome occupancy of mRNA at the beginning of the open reading frame increase^25, 34, 35^, and that de-repression of translation initiation *(gcn2Δ)* is insufficient to recover protein synthesis^23,25,26,34,35^. Temporary translation repression is an essential part of environmental stress response,^24^ and our results consistently suggested that K63 ubiquitin affects ribosome activity *post* initiation.

Our results also indicated that this role resulted from differential effects of K63 ubiquitin on translation factors and their interaction with the ribosome (Figure 5). While a detailed mechanism remained to be revealed, K63 ubiquitin was essential for proper association of these factors with the ribosome (Figure 4 and 5). Dysregulated binding of elongation factors (e.g. eEF2) could impact protein production. Dysregulated binding of initiation factors (e.g. eIF3) could affect mRNA recruitment, 40S-60S association, or 40S scanning, acting as fail-safe mechanisms in response to stress.

Almost two decades after the initial discovery of K63 ubiquitin modification of a single ribosomal protein, Rpl28 (uL15)^36^, only few large-scale studies of ribosome ubiquitination exist to date. One such recent study described ubiquitinated sites at the 40S subunit during the unfolded protein response induced by stress of the endoplasmic reticulum^10^. The residues were modified primarily by mono-ubiquitin or K48 di-ubiquitin. Other work highlighted the role of the ZNF598 ubiquitin ligase which is a *HEL2* homolog in yeast and functions in quality control to resolve stalled ribosomes^11, 12, 37^. Given the large number of ubiquitin enzymes encoded in the genome, the diversity of ubiquitin chains, and the number of modifiable lysine residues in the ribosome, the putative regulatory network governing ubiquitination is extremely complex, and perhaps as intricate as that of histone modifications^38^. We would argue that tools and data as presented here provide important steps towards elucidating this role.

## Supporting Information

The Supporting Information is available free of charge Supplementary Figures S1–S6 (PDF)

Supplementary Table S1 – SILAC Quantification of ubiquitinated proteins (.XLSX)

Supplementary Table S2 – Ribosome profiling protein quantification (.XLSX)

Supplementary Table S3 – Yeast strains used in this study (.XLSX)

## Acknowledgements

We thank D. Finley for providing yeast strains, J.R. Cussiol for the yeast deletion templates and support in yeast genetics. We are indebted to Evan Baugh for the assistance with the structural mapping.

The K63 ubiquitin mass spectrometric experiments were conducted in the NYU Proteomics Resource Center (Dr. Ueberheide), which is in part supported by the NYU School of Medicine.

We thank M. Beck, P. Kastritis, M. Gorospe, and H. D. Ryoo for the scientific discussions and comments on the manuscript.

This work was supported in part by the US National Institutes of Health K99/R00 award ES025835 (GMS). CV acknowledges funding by the NIH/NIGMS (R01 GM113237 and 1R35GM127089-01), and the Zegar Family Foundation Fund for Genomics Research at New York University.

The authors declare no financial interest. The funders had no role in study design, data collection and analysis, decision to publish, or preparation of the manuscript.

## Author Information

### Author Contributions

G.M.S. designed the project. S.B and G.M.S. performed the experiments. C.V. and G.M.S. analyzed the data. C.V. and G.M.S. wrote the manuscript. All authors discussed the results and commented on the manuscript.

The mass spectrometry proteomics data have been deposited to the ProteomeXchange Consortium via the PRIDE partner repository^39^ with the dataset identifier PXD004650.

## Supplementary Figure Legends

**Supplementary Figure 1.**
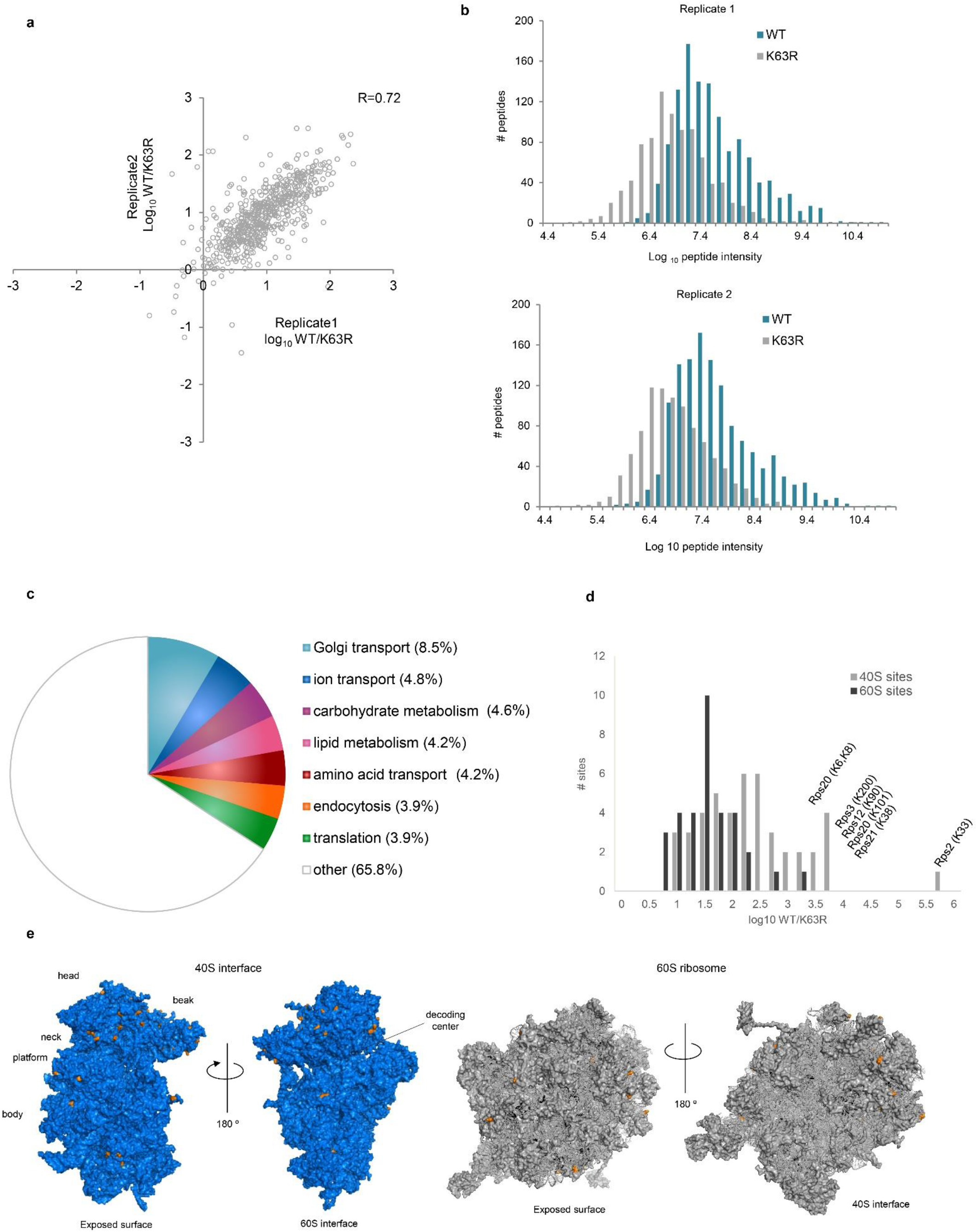
K63 ubiquitinomics in response to oxidative stress. **(a)** Scatter plot of SILAC log_10_ WT/K63R ratio of K63 GG peptide intensity from two independent replicates. **(b)** Frequency distribution of log_10_ intensity from WT (teal) and K63R (gray) for two independent replicates. **(c)** Pie chart of GO annotations (SGD – biological process) for K63 target proteins. **(d)** Frequency distribution of SILAC log_10_ WT/K63R ratio of K63 GG peptide intensity from 40S ribosomal proteins (gray) and 60S ribosomal proteins (black). Highly abundant sites are highlighted. **(e)** Mapping of the K63 ubiquitinated sites (orange) to the 3D surface structure of the yeast ribosome (PDB id:4v88)^20^. 40S subunit is presented in blue and 60S subunit is presented in gray. WT, wild-type GMS280 yeast strain. K63R, ubiquitin K63R mutant GMS413 yeast strain.

**Supplementary Figure 2.**
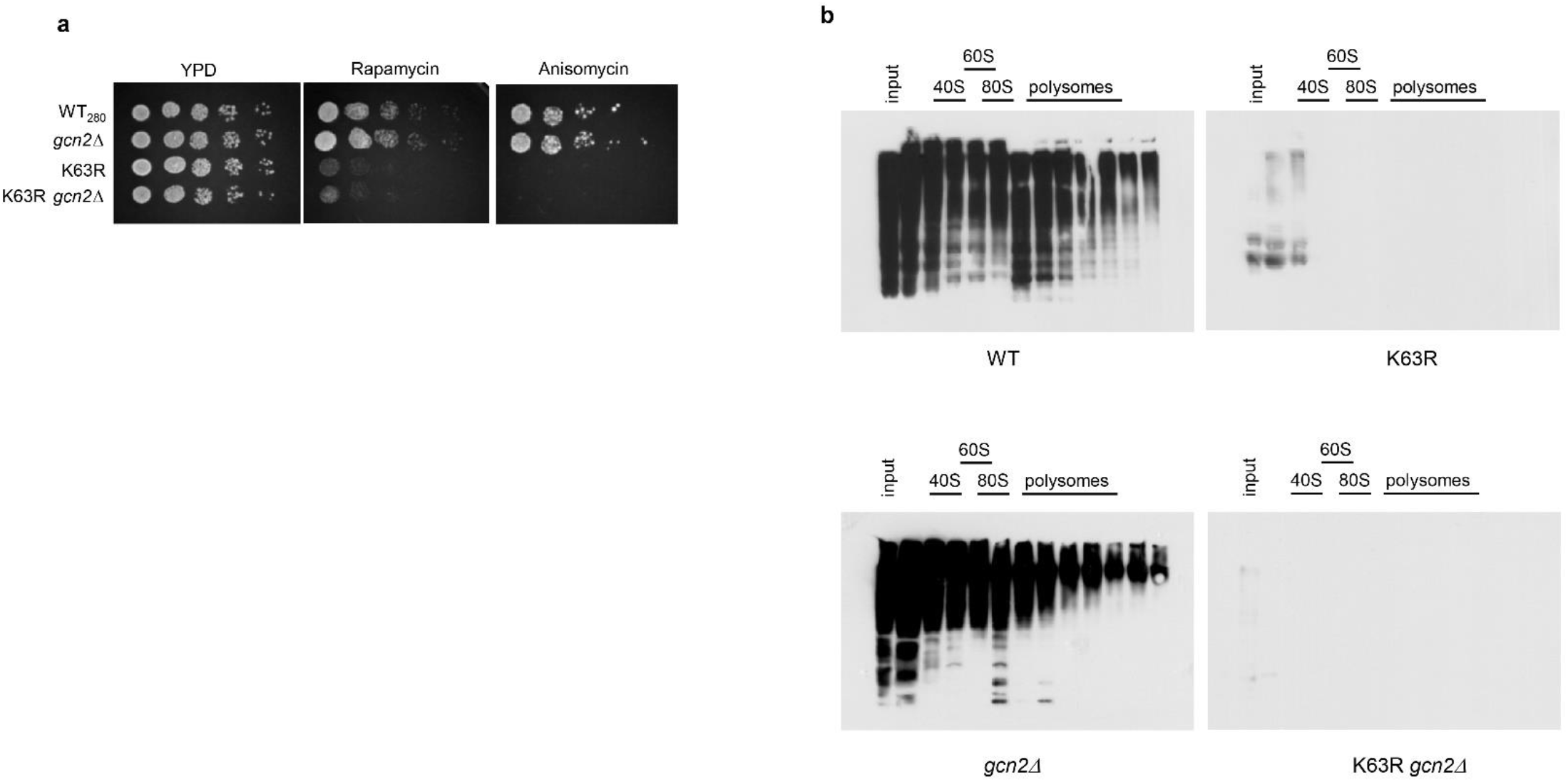
Deletion of *GCN2* does not reverse K63R mutant sensitivity to translation inhibitors. **(a)** Serial dilution assay in the presence of 50 ng/ml rapamycin or 20 μg/ml anisomycin in rich YPD medium. **(b)**. Anti-K63 ubiquitin western blot of polysome fractions from the designated cells treated with 0.6mM H_2_O_2_ for 30 min. Accompanying polysome profiles are presented in Figure 3A-D.

**Supplementary Figure 3.**
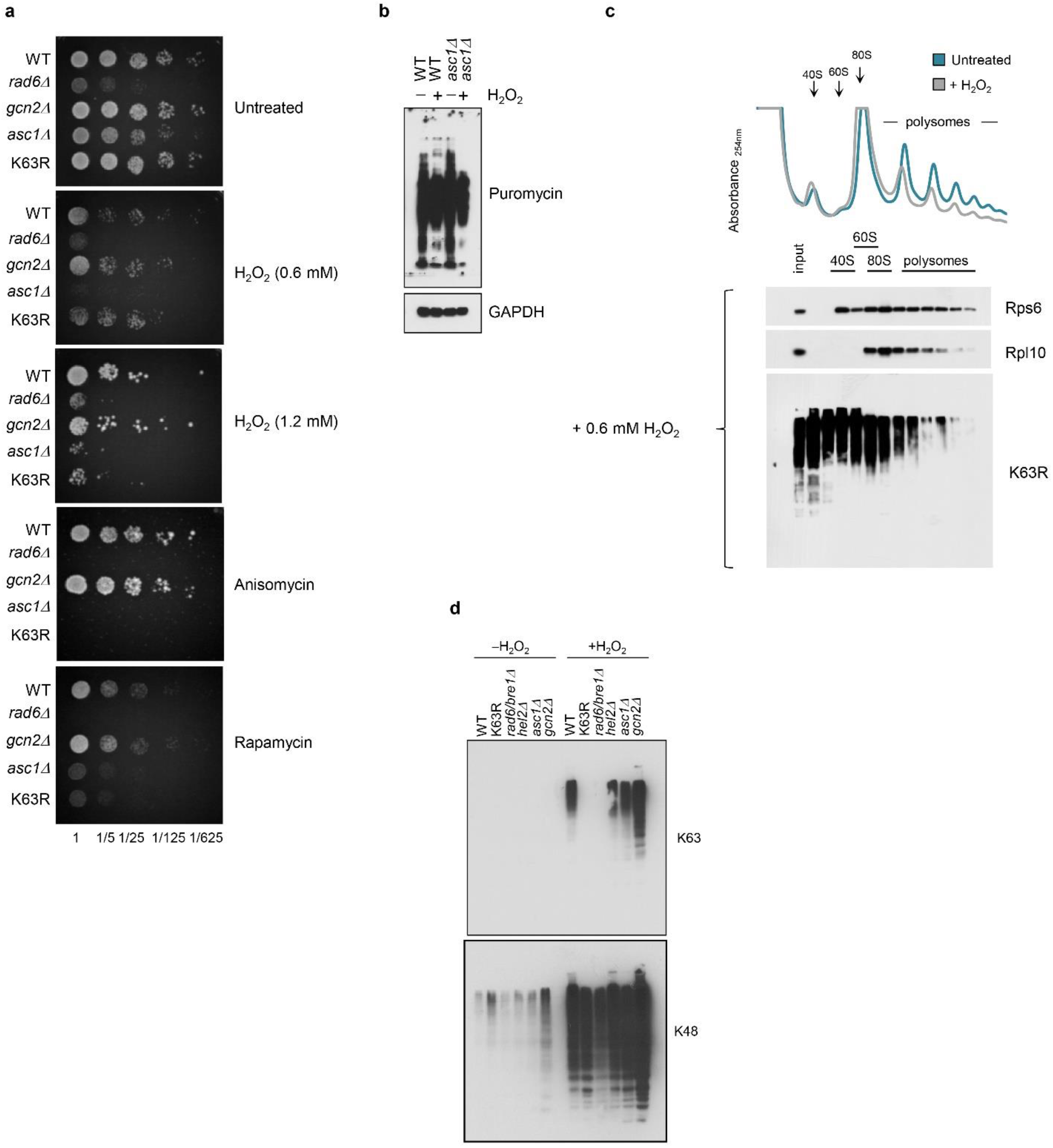
Asc1 (RACK1) is not involved in the K63 ubiquitin redox pathway. **(a)** Serial dilution assay in the presence of 0.6mM or 1.2 mM H_2_O_2_, 50 ng/ml rapamycin or 20 μg/ml anisomycin in rich YPD medium **(b)** Western blot anti-puromycin was used to evaluate *de novo* protein synthesis. Cells were treated with H_2_O_2_ for 15 min prior to 30 min treatment with 0.9 mM puromycin. Anti-GAPDH was used as loading control. **(c)** Sucrose sedimentation profiles of polysome from *asc1 Δ* strain in the presence or absence of H_2_O_2_. Western blot of sedimentation fractions was performed for the 60S large subunit protein Rpl10, the 40S small subunit protein Rps6, and K63 ubiquitin. **(d)** Hel2 (E3 ubiquitin ligase) is not responsible for K63 ubiquitination in response to oxidative stress. Western blot anti-K63 and K48 ubiquitin was performed in the presence or absence of oxidative stress induced by 30 min of 0.6 mM H_2_O_2_ treatment.

**Supplementary Figure 4.**
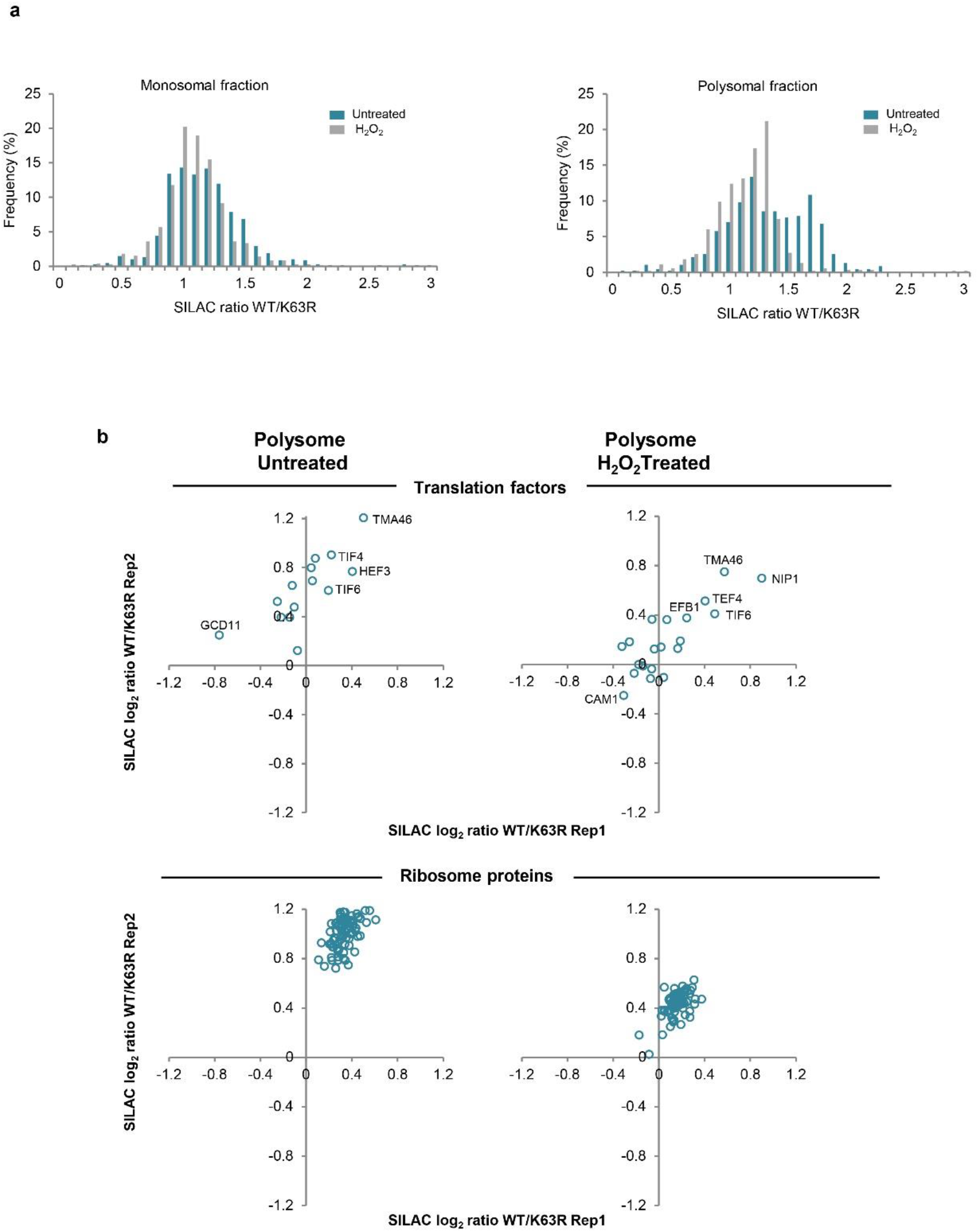
Ribosome proteomics of the polysome fraction. Frequency distribution of SILAC log_10_ WT/K63R ratio of protein intensity in the absence (teal) or the presence (gray) of H_2_O_2_ from the **(a)** monosome and polysome fraction. **(b)** Scatter plot of SILAC WT/K63R ratios of protein intensity from two independent replicates in the polysome fraction. *(top)* SILAC ratios of translation factors, *(bottom)* SILAC ratios of ribosomal proteins.

**Supplementary Figure 5.**
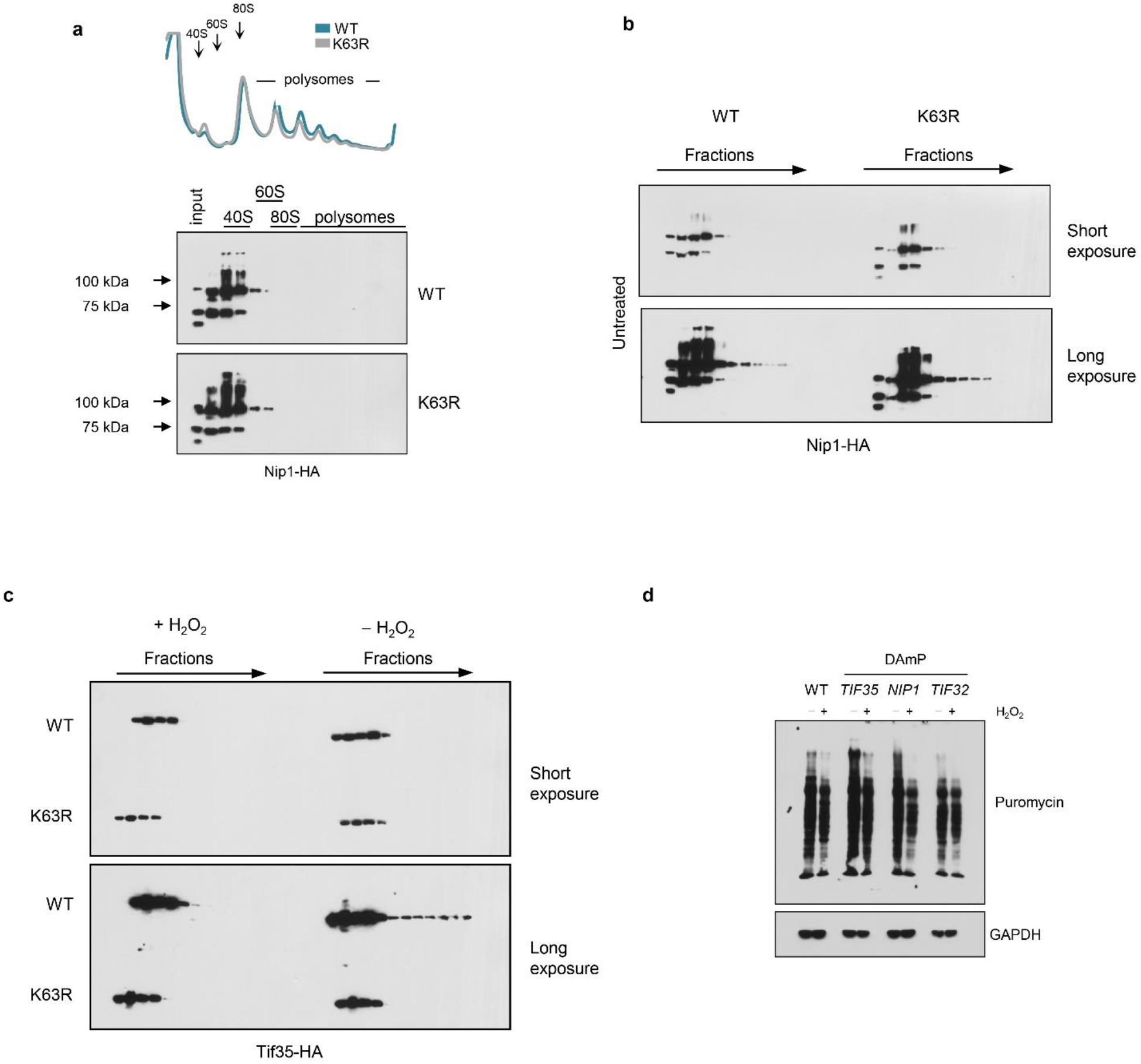
Nip1 is not enriched in the wild-type ribosomes. **(a)** Sucrose sedimentation profiles of polysome from wild-type and K63R mutant bearing HA-tagged Nip1 in the absence of H_2_O_2_. Blotting against HA shows most of the Nip1 protein bound to the 40S fraction. Results are not dependent on blot exposure. **(b)** Nip1-HA blots of WT and K63R were exposed for different amounts of time. **(c)** Tif35-HA blots for wild-type and K63R mutant strain were exposed for different amounts of time and show differential binding to wild-type ribosomes. **(d)** Western blot against puromycin was used to evaluate *de novo* protein synthesis from wild-type, and different yeast strains from the DAmP collection. Anti GAPDH blot was used as loading control.

**Supplementary Figure 6.**
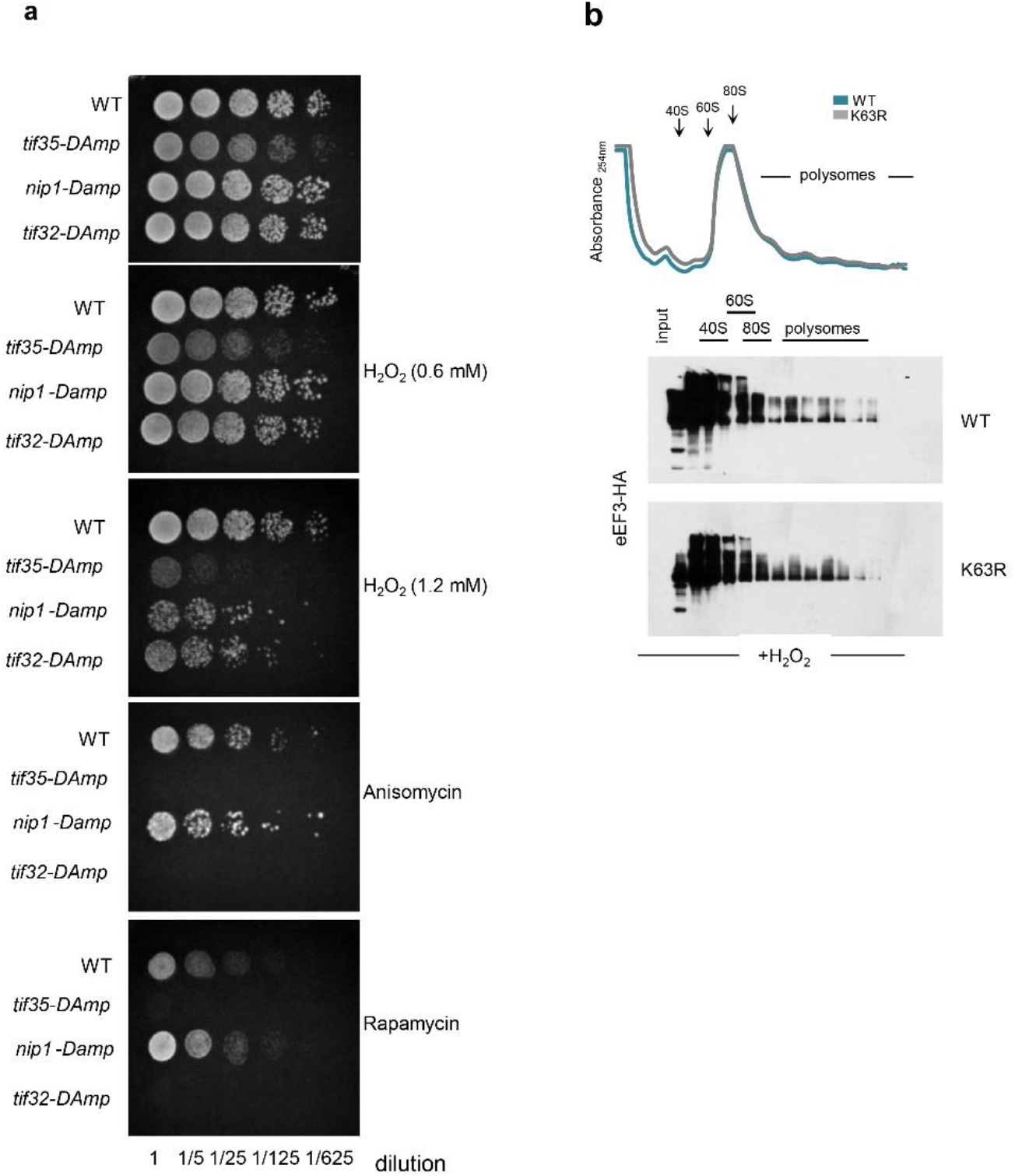
Destabilization of eIF3 subunit transcripts sensitized cells to oxidative stress and translation inhibitors. **(a)** eIF3 strains from the DAmP collection were subjected to serial dilution assay in the presence of 0.6mM or 1.2 mM H_2_O_2_, 50 ng/ml rapamycin or 20 μg/ml anisomycin in rich YPD medium. **(b)** eEF3 binding to ribosomes is not dependent on K63 ubiquitin.

